# Overexpression of TPM4 promotes cancer progression through EMT in anaplastic thyroid cancer

**DOI:** 10.1101/2025.11.22.689948

**Authors:** Hangjie Ying, Yahui Zhou, Xiaoming Jiang, Wei Cui, Shanshan Chen, Xiaofang Yao, Jianguo Feng, Dong Guo, Xu Qian

## Abstract

Anaplastic thyroid cancer (ATC) is an undifferentiated tumour originating from thyroid follicular epithelium and represents the most malignant subtype of thyroid cancer. However, the mechanisms underlying ATC development are not fully understood. This study aimed to investigate the mechanisms driving ATC invasion and metastasis to provide novel insights into its aggressive behavior. Through an integrated bioinformatics analysis of the GEO database, we identified Tropomyosin 4 (TPM4) as a differentially expressed gene in ATC. Bioinformatic correlation and pathway enrichment analyses suggested that TPM4 expression is potentially linked to tumor cell invasion and lymph node metastasis. We showed that TPM4 was highly expressed in both ATC tissues and cell lines, and its expression levels were positively correlated with enhanced cell migration and invasion. Mechanistic studies indicated that TPM4 likely enhances tumour cell migration and invasion by facilitating cytoskeletal remodeling and promoting the epithelial-mesenchymal transition (EMT). Our findings suggest that TPM4 represents a promising diagnostic and therapeutic biomarker for ATC.

## Introduction

Anaplastic thyroid cancer (ATC) is a rare thyroid tumour with a highly aggressive malignancy, accounting for approximately 1-2% of all thyroid cancers. It typically arises from pre-existing differentiated thyroid cancers through the accumulation of specific genetic alterations^1,2^. ATC manifests as a rapidly growing, infiltrative neck mass and is invariably classified as stage IV disease at diagnosis due to its extreme aggressiveness^3^.The disease-specific mortality rate approaches 100%, with median survival of only 3 to 6 months^4,5^. Approximately 20-50% of ATC patients develop distant metastases, most commonly to the lungs, bones, and brain^6,7^. Immunotherapy combining checkpoint inhibitors with anti-angiogenic agents or BRAF/MEK inhibitors has demonstrated significant efficacy in some ATC patients, even achieving long-term survival^8-10^. Despite these advances, the high metastasis rate and frequent emergence of therapeutic resistance remain formidable challenges^11^. Therefore, the identification of novel biomarkers for the early diagnosis and improved treatment of ATC is urgently needed.

The tropomyosin family (TPM) comprises a group of proteins closely associated with the cytoskeleton and actin, encompassing four major genes: TPM1, TPM2, TPM3, and TPM4 ^12^. TPM4 is an actin-binding protein that provides structural stability to actin filaments and regulates cytoskeletal function. By modulating cell-extracellular matrix (ECM) interactions, the cytoskeleton plays a critical role in tumour metastasis^13^. Extensive research indicates that abnormal expression of TPM4 is associated with the development and progression of multiple cancers, including gastric^14^, colorectal^15^, breast^16^, lung, ovarian^17^, oesophageal^18^, hepatocellular^19^, oral squamous cell ^20^, cervical^21^, and prostate ^22^. In papillary thyroid carcinoma (PTC), hypoxia-induced tumour microenvironments promote lymph node metastasis through TPM4-mediated epithelial-mesenchymal transition (EMT) signalling activation^23^. However, the expression and biological role of TPM4 in ATC remain unexplored.

Previous studies suggest TPM4 might also regulate the proliferation-differentiation balance, demonstrating its robust expression in proliferating and undifferentiated tumour cell lines (Caco-2) that can be induced to mature, and suppression during growth arrest and maturation^24^. It is recognized that both ATC and PTC originate from thyroid follicular epithelial cells, yet they exhibit entirely distinct differentiation levels, malignant potential, and prognostics^25^. Based on these studies, we hypothesize that TPM4 dysregulation may serve as a molecular event predisposing thyroid tissue to tumourigenesis, sustaining proliferation and maintaining an undifferentiated state. Its overexpression may represent an early form of genetic damage contributing to the development and malignant proliferation of ATC. It remains to be investigated whether TPM4 expression differs markedly in ATC compared to other thyroid cancers, how it contributes to tumor proliferation, metastasis, and poorer prognosis, and what the underlying mechanisms are. Our study, through bioinformatics analysis and experimental validation, aims to elucidate the roles of TPM4 in ATC cell proliferation, migration, and invasion, and to explore its potential value in ATC clinical prognosis.

## Methods

### Datasets download and RNA-seq analysis

This study utili sed four publicly available expression datasets from the Gene Expression Omnibus (GEO) database (GSE29265, GSE33630, GSE65144, and GSE76039), all generated from the same human gene expression microarray platform (GPL570, Affymetrix Human Genome U133 Plus 2.0 Microarray). GSE29265 contains 9 ATC samples, 20 PTC samples, and 20 normal thyroid tissue samples^26^. GSE33630 comprises 11 ATC samples, 49 PTC samples, and 45 normal thyroid tissue samples^27^. GSE65144 comprises 12 ATC samples and 12 normal thyroid tissue samples^28^. GSE76039 contains 20 ATC samples^29^. All datasets are publicly available and accessible via the website (https://www.ncbi.nlm.nih.gov/geo/). Data acquisition and utilisation adhere to the principles and guidelines of the GEO database, in accordance with its data access policies. Data merging and analysis were performed using R software (version 4.4.0). The ‘Bioconductor’ package was employed to merge datasets, utilising the removeBatchEffect function for batch effect correction. Principal Component Analysis (PCA) was applied to evaluate the efficacy of batch effect removal and to assess differences or distribution patterns between samples. The threshold for DEGs was set at an adjusted p-value (adj.P.Val) < 0.05 and absolute |logFC| > 1. Based on the identified DEGs, heatmaps were generated using the ‘pheatmap’ package, and volcano plots were created with the ‘ggplot2’ package to display gene logFC and statistical significance. Functional annotation and pathway enrichment analysis were performed in R using the ‘clusterProfiler’, ‘org.Hs.eg.db’, and ‘enrichplot’ packages^30^, including Gene Ontology (GO) and Kyoto Encyclopedia of Genes and Genomes (KEGG) enrichment ^31^. The GO enrichment analysis encompassed the categories of molecular function (MF), biological process (BP), and cellular component (CC).

### Clinical data analysis

Gene expression samples for thyroid cancer, alongside patient lymph node metastasis status and survival information, were downloaded from the cBioPortal-TCGA database (https://www.cbioportal.org/)^32-35^. Thyroid cancer patients were categorised into “high” and “low” TPM4 expression groups based on median TPM4 expression levels. Analysis was conducted using R software (v 4.4.0, The R Foundation for Statistical Computing, Vienna, Austria). Samples were categorised based on lymph node metastasis status: no metastasis (N0), metastasis present (N1), and normal thyroid tissue samples (from the cBioPortal datasets). Comparisons were performed using the Kruskal-Wallis H test, with P < 0.05 indicating statistical significance.

### Immunohistochemical staining

Immunohistochemical (IHC) staining was performed on formalin-fixed and paraffin-embedded (FFPE) sections of ATC and adjacent normal thyroid tissue, following a previously established protocol^36^. A mouse-derived primary antibody against TPM4 (Cat. No. Ag6947, Proteintech, China) was used for staining. Images were captured using an Olympus SLIDEVIEW VS200 digital slide scanner and analysed with Olympus Olyvia software (v3.3).

### Cell lines and cell culture

The human ATC cell lines KHM-5M, KMH-2, C643, and CAL-62 were obtained from the Chinese Academy of Sciences Cell Bank (China). All cell lines were maintained in Dulbecco’s Modified Eagle Medium (DMEM) or Roswell Park Memorial Institute (RPMI) 1640 medium, supplemented with 10% fetal bovine serum (FBS), at 37°C in a humidified atmosphere containing 5% CO_2_. Cells were harvested during the logarithmic growth phase for subsequent experiments.

### Western blot analysis of TPM4 expression

After cells were collected, they were lysed thoroughly with 200 μL of lysis buffer (containing RIPA, broad-spectrum protease inhibitors, phosphatase inhibitors, and PMSF in a ratio of 100:2:2:1). The protein lysate was mixed with 5 × loading buffer at a 4:1 ratio and denatured by heating at 95°C for 10 minutes. Subsequently, the proteins were separated by SDS-PAGE and transferred to a PVDF membrane. The membrane was blocked with 5% skimmed milk for two hours at room temperature and then incubated with the primary antibody against TPM4 overnight at 4°C. After three washes with TBST buffer, the membrane was incubated with the secondary antibody for one hour at room temperature. Protein signals were detected using a chemiluminescence imaging system.

### Transwell invasion assay

Cell invasion was assessed using 24-well Transwell chambers. The upper chamber was coated with a layer of Matrigel (diluted 1:8 in serum-free DMEM medium). Cells were resuspended at a density of 1 × 10^5^ cells per well and seeded into the upper chamber. The lower chamber was filled with DMEM medium supplemented with 10% FBS as a chemoattractant. The plates were then incubated for 48 hours at 37°C in a 5% CO_2_ atmosphere. After incubation, the cells on the upper surface of the membrane were carefully removed. The cells that had invaded through the Matrigel and membrane pores to the lower surface were fixed with 4% paraformaldehyde, stained with 0.1% crystal violet, and quantified by counting under a microscope.

### CCK-8 assay

C643 and CAL-62 cells were seeded into 96-well plates at a density of 2 × 10^4^ cells per well in 100 μL of culture medium. The plates were incubated at 37°C in a 5% CO_2_ incubator. At 24, 48, and 72 hours after seeding, 10 μL of CCK-8 solution was added to each well, followed by incubation for another two hours. The optical density (OD) at 450 nm was then measured for each well using a microplate reader.

### Scratch assay

C643 and CAL-62 cells with TPM4 knockdown and the corresponding control cells were seeded into six-well plates and cultured until they reached approximately 90% confluence, forming a confluent monolayer. A straight scratch was created in the monolayer using a sterile 10 μL pipette tip. The dislodged cells were washed away with PBS, and fresh medium supplemented with 1% FBS was added. Images of the scratch wounds were captured under a microscope immediately (0 h) and 24 hours after wounding. The relative migration distance was compared between groups by measuring the change in the scratch width.

### Enzyme-Linked Immunosorbent Assay

Plasma samples from patients with pathologically diagnosed ATC, PTC, and healthy controls were collected (Supplementary Table 1).

The pre-coated 96-well plate from the commercial human TPM4 ELISA kit (Cat. No. ml106375, Mlbio, China) was used. Briefly, the plate was equilibrated to room temperature. Diluted standards and plasma samples were then added to the designated wells and incubated for two hours at 37°C. After washing, a horseradish peroxidase (HRP)-conjugated detection antibody was added and incubated for one hour at 37°C. Following another wash, tetramethylbenzidine (TMB) substrate was added and incubated in the dark at room temperature for 15-30 minutes for color development. The reaction was stopped by adding the stop solution, and the absorbance was immediately measured at 450 nm using a microplate reader. The concentration of TPM4 in each sample was interpolated from the standard curve. **Quantitative and statistical analysis**

All quantitative data were statistically analysed using ImageJ software. Statistical significance was assessed using the two-tailed Student’s *t*-test or Welch’s *t*-test. Differences with p < 0.05 were considered statistically significant. All measurements are presented as mean ± SEM.

## Results

### Bioinformatics analysis and experimental validation identify TPM4 as a potential biomarker in ATC

We integrated four GEO datasets (GSE29265, GSE33630, GSE65144, and GSE76039) from the GPL570 platform, comprising 52 ATC, 69 PTC, and 77 normal thyroid tissue samples after batch-effect correction (Fig. 1A). Differential expression analysis identified 2,318 genes in ATC versus normal tissue (1,138 up, 1,180 down), 740 in ATC versus PTC (260 up, 480 down), and 2,313 in PTC versus normal tissue (1,216 up, 1,097 down) (Supplementary Fig. 1). Among these, 222 genes exhibited differential expression across all three comparisons, with 83 commonly upregulated and 139 commonly downregulated.

**Figure 1.**
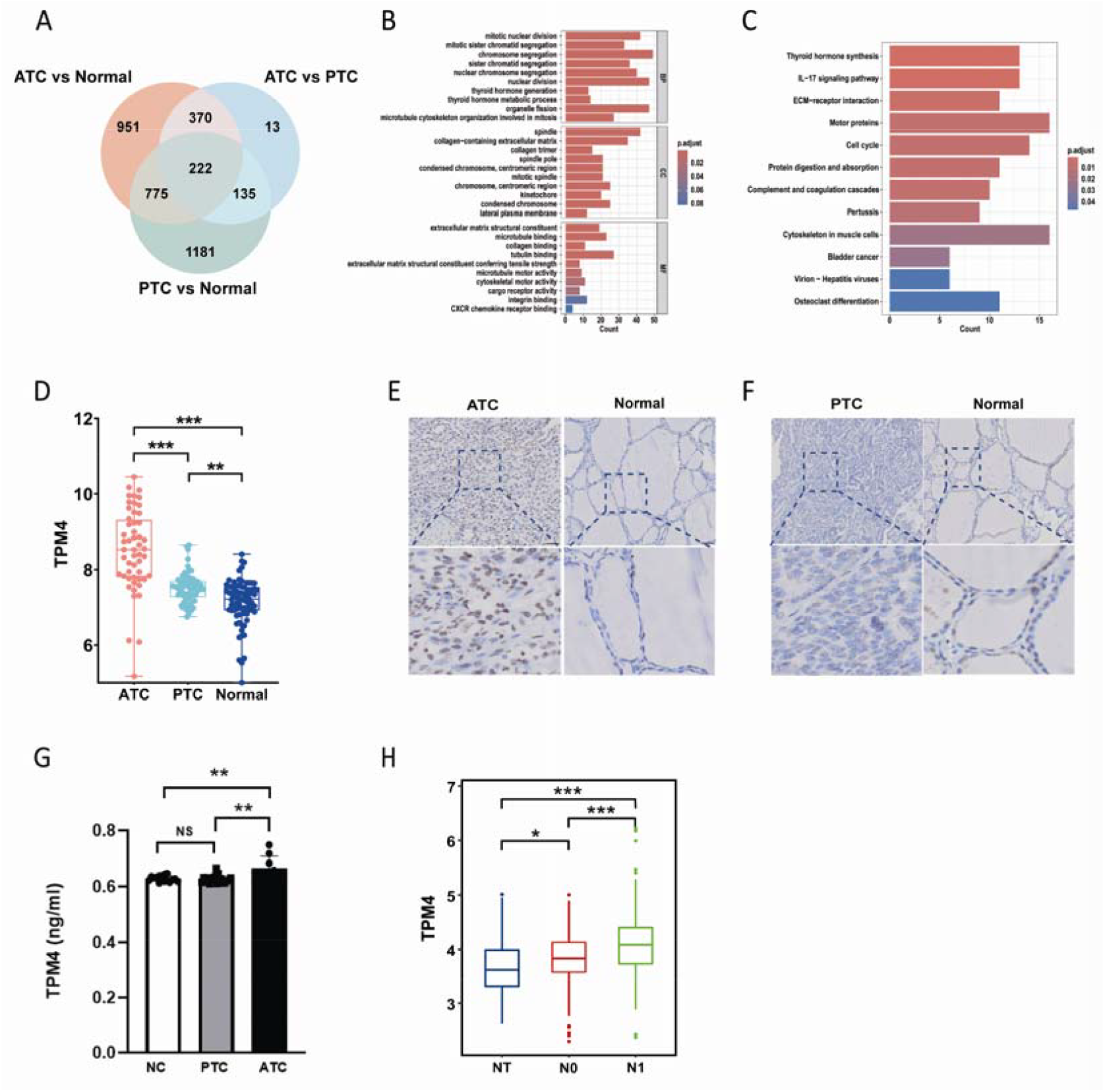
Bioinformatics analysis and experimental validation identify TPM4 as a potential biomarker in ATC. (A) Venn diagram of the 222 common differentially expressed genes (DEGs) across three comparisons: ATC versus normal tissues, ATC versus PTC, and PTC versus normal tissues. (B) GO enrichment analysis of the common DEGs. (C) KEGG pathway enrichment analysis of the common DEGs. (D) The box plot shows the expression of TPM4 in the three groups: ATC, PTC, and normal. (E,F) TPM4 is highly expressed in ATC and PTC. (G) Plasma TPM4 protein levels in ATC, PTC and normal samples (NC) was detected by ELISA (**p<0.01, by one-way ANOVA with post-hoc test). (H) TPM4 expression levels across NT, N0 (no lymph node metastasis), and N1 (lymph node metastasis present) groups (*p <0.05, ***p<0.001, by one-way ANOVA with post-hoc test).

GO enrichment analysis of these 222 shared genes revealed their involvement in biological processes including mitotic cell division, extracellular matrix organization, and thyroid hormone metabolism (Fig. 1B). KEGG pathway analysis identified enriched pathways including complement and coagulation cascades, IL-17 signaling pathways, extracellular matrix-receptor interactions, and cytokine-cytokine receptor interactions (Fig. 1C).

Furthermore, the dysregulated genes in ATC tissues were prominently associated with cell cycle progression and cancer pathways, suggesting the heightened proliferative activity of ATC cells.

Previous studies have identified hypoxia-induced upregulation of TPM4 activating the EMT signalling pathway to promote lymph node metastasis in PTC^23^. In our study, RNA-seq data indicate that TPM4 transcriptional levels are significantly higher in the more malignant ATC subtype compared to PTC and healthy thyroid tissue (Fig. 1D). The expression profile of TPM4 in ATC showed considerable heterogeneity, characterized by a wider distribution and more extreme outliers compared to other groups. Concurrently, ELISA and IHC (Fig. 1E-G) results consistently demonstrated that TPM4 is highly expressed in ATC, with no apparent upregulation observed in PTC. This elevated expression, coupled with its positive correlation with lymph node metastasis, aligns with the highly metastatic and aggressive nature of ATC. Furthermore, TPM4 expression levels showed a positive correlation with lymph node metastasis in thyroid carcinoma (Fig. 1H).

### TPM4 promotes proliferation, migration, and invasion of ATC cells

Based on Western blot analysis which identified C643 and CAL-62 as the ATC cell lines with the highest TPM4 expression among those tested (KHM-5M, KMH-2, C643, and CAL-62) (Fig. 2A), we proceeded to knock down TPM4 in these two cell models. Knockdown of TPM4 was performed using three distinct shRNAs (sh-1, sh-2, and sh-3), all of which significantly reduced TPM4 protein levels. Among them, sh-3 yielded the most potent knockdown in both cell lines (Fig. 2B) and was therefore used in all subsequent experiments.

**Figure 2.**
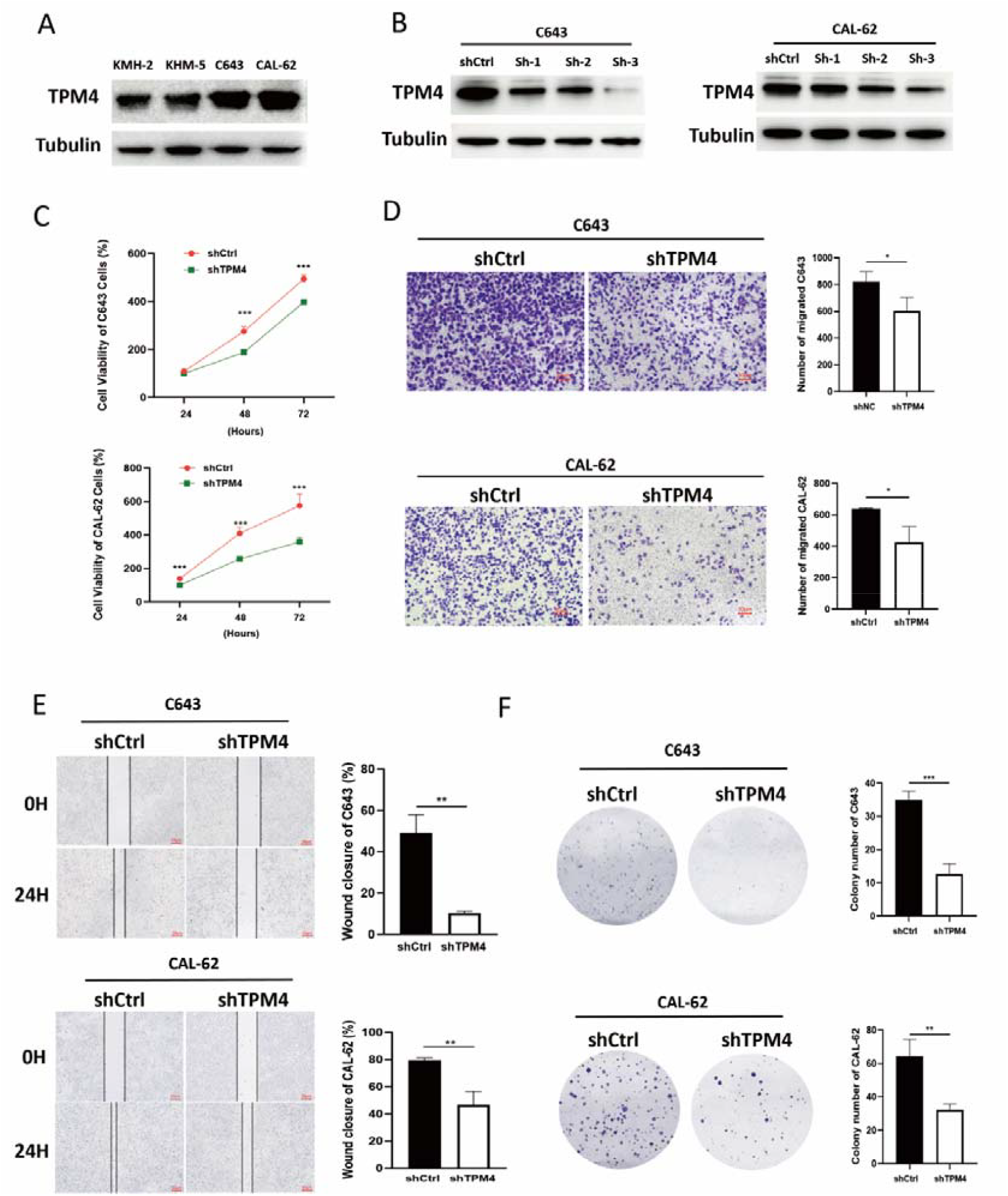
TPM4 promotes proliferation, migration, and invasion of ATC cells. (A) Western blot analysis of TPM4 expression in various ATC cell lines. (B) Western blot analysis of TPM4 knockdown efficiency in C643 and CAL-62 cells after shRNA infection. (C) Effects of TPM4 knockdown on the proliferation of ATC cells assessed by CCK-8 assay. (D) Comparison of invasive abilities between control and TPM4-knockdown ATC cells. (E) Wound-healing assay assessing migration of C643 and CAL-62 cells with or without TPM4 depletion in 1% FBS medium. (F) Colony formation assay of C643 and CAL-62 cells with or without TPM4 depletion after two weeks of culture. All quantitative data were statistically analyzed using Image J software. Statistical analysis was performed using GraphPad Prism 8.0 with Student’s *t*-test (*p < 0.05, **p < 0.01, ***p < 0.001).

The efficient knockdown of TPM4 significantly suppressed cell proliferation, as demonstrated by reduced OD values in CCK-8 assays at 48 and 72 hours (Fig. 2C) and a marked reduction in colony formation capacity (Fig. 2F). Furthermore, TPM4 depletion impaired metastatic capabilities, leading to a significant decrease in cell invasion (Fig. 2D) and migration (Fig. 2E). Collectively, these results demonstrate that TPM4 plays a crucial role in promoting proliferation and invasion of ATC cells.

### TPM4 is required for epithelial to mesenchymal transition

TPM4 knockdown induced pronounced morphological alterations in both C643 and CAL-62 cells, which became more flattened and exhibited weakened intercellular junctions, suggesting a critical role for TPM4 in maintaining cellular architecture (Fig. 3A). Consistent with a loss of mesenchymal features, immunoblotting analysis showed that TPM4 knockdown decreased the expression of mesenchymal markers (N-cadherin and Vimentin) while increasing the expression of the epithelial marker E-cadherin (Fig. 3B). Collectively, these results indicate that TPM4 promotes ATC cell invasiveness by regulating the expression of key EMT-associated proteins.

**Figure 3.**
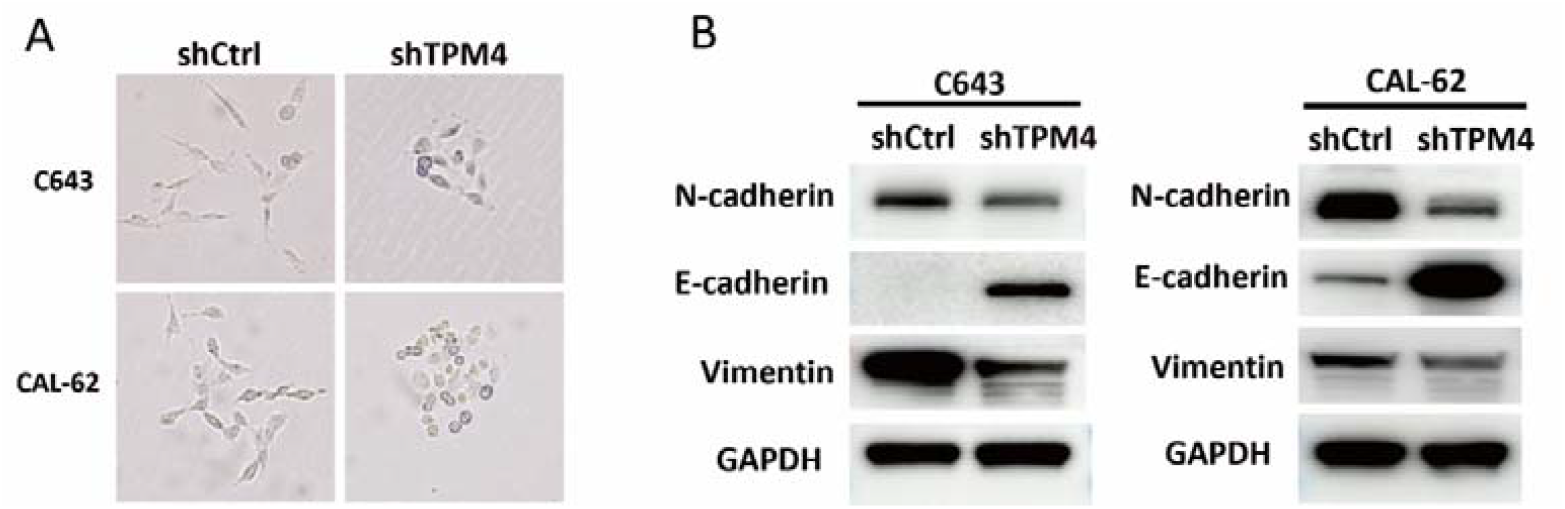
TPM4 is required for epithelial to mesenchymal transition. (A) Representative phase-contrast images showing morphological changes in C643 and CAL-62 cells following TPM4 knockdown. Original magnification, × 200. (B) Immunoblotting analyses with the indicated antibodies were performed.

## Discussion

During tumourigenesis and progression, cytoskeletal remodelling plays a pivotal role. Cell migration relies upon coordinated actin cytoskeletal organisation. Cell migration and invasion are central to morphogenesis and tumour metastasis, with coordinated interactions of the cytoskeleton being indispensable in this process. Actin, microtubules, and intermediate filaments maintain continuous mutual interactions, collectively propelling the migration process through their dynamically coordinated changes^37,38^. TPM4 is an actin-associated protein that influences cell adhesion to the ECM by cross-linking with α-actin to form actin stress fibres, thereby stimulating cell migration in tumours^15^. Furthermore, TPM4 plays a pivotal role in the initiation and progression of multiple cancers, being intrinsically linked to biological processes such as cell motility, proliferation, and metastasis^13,17,39^. However, investigations into the role and mechanisms of the TPM4 gene in the development and progression of ATC remain insufficient.

Our integrated bioinformatics and experimental approach (IHC, ELISA) consistently demonstrated that TPM4 is specifically upregulated at the transcriptomic and protein levels in ATC, but not in PTC, compared to normal tissue. This suggests that elevated TPM4 expression may contribute to molecular mechanisms underpinning increased tumour malignancy, potentially through affecting cell differentiation. Previous studies identified TPM4 upregulation activating the epithelial-mesenchymal transition (EMT) signalling pathway to promote highly metastatic PTCs. Clinical data analysis revealed a significant positive correlation between TPM4 expression levels and lymph node metastasis in TC patients, consistent with prior findings^23^. To further investigate whether TPM4 modulates proliferation, invasion, and migration in ATC cells, we knocked down TPM4 expression in two major ATC cell lines (C643 and CAL-62). Results demonstrated TPM4 knockdown markedly reduced proliferation, invasion, and migration. This indicates TPM4 promotes malignant proliferation in ATC cells.

EMT is recognized as a primary driver of cancer progression from initiation to metastasis. It constitutes a biological process whereby cells undergo a shift from an epithelial phenotype to a mesenchymal phenotype, characterised by morphological alterations, changes in the cytoskeleton, loss of intercellular adhesion, and the acquisition of invasive and metastatic capabilities^40,41^. Our Western blot analysis revealed that TPM4 knockdown in the ATC cell line resulted in increased E-cadherin expression alongside decreased N-cadherin and vimentin expression. These findings indicate that TPM4 expression in ATC cells promotes the EMT process, thereby enhancing the invasive and migratory capabilities of these cells.

In summary, our research indicates that TPM4 plays a significant role in the initiation, progression, and invasive metastasis of ATC. It enhances tumour cell migration and invasive capacity by regulating cytoskeletal remodelling and promoting EMT. TPM4 holds potential as a biomarker for ATC diagnosis and treatment.

## Acknowledgments

This study was supported by grants from the National Natural Science Foundation of China (82272406, X.Q., 82522060, 82372816, D.G.).

## Author contributions

X.Q. conceived and designed the study; H.Y, Y.Z., X.J., W.C., S.C. and X.Y. performed the experiments; D.G. and J.F. provided reagents, technical support, and conceptual advice; H.Y. and Y.Z. wrote the paper; all authors commented on the paper.

---

**Supplementary Figure 1.**
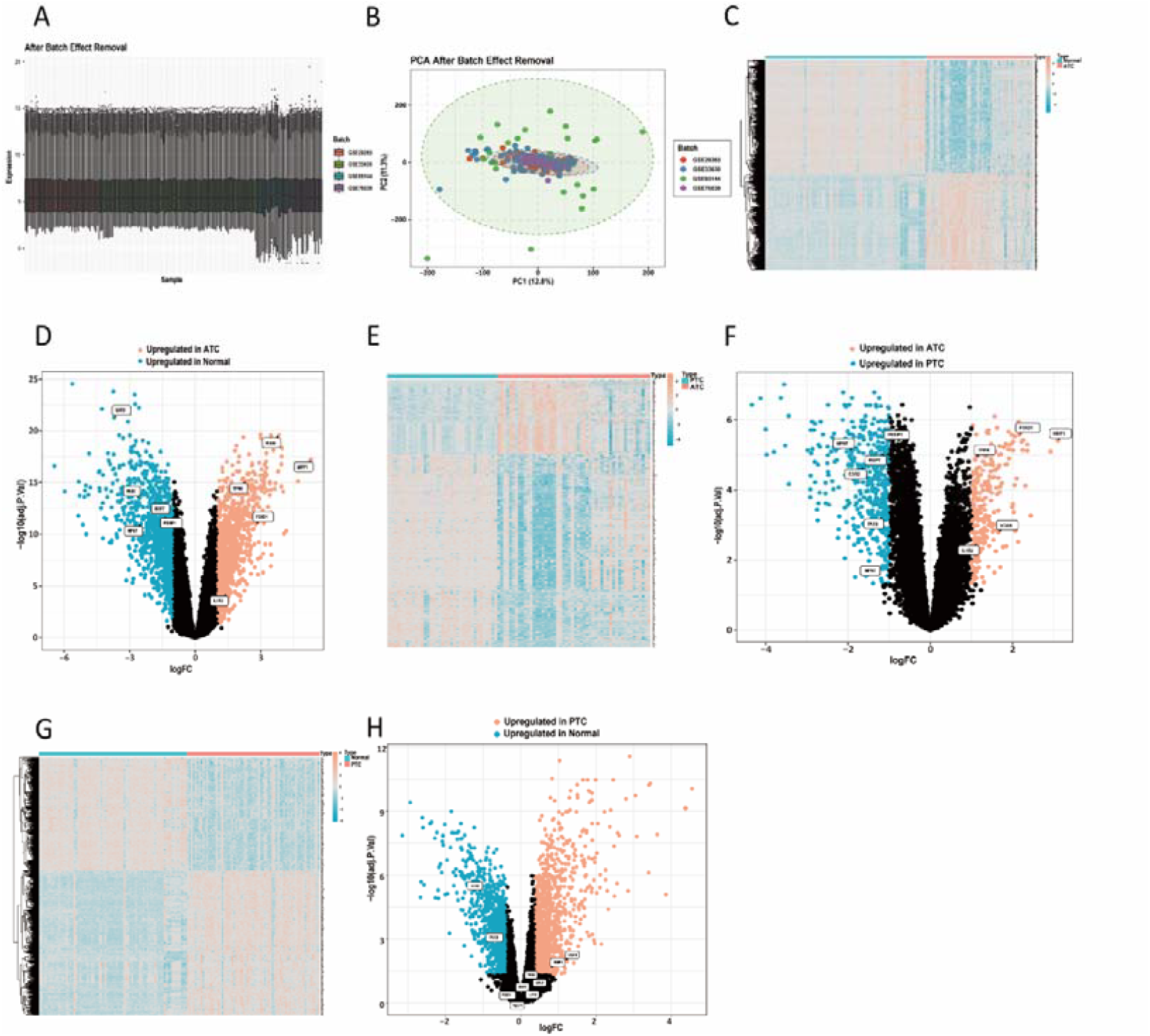
(A) Box plots showing normalized gene expression distributions across all samples from the four integrated datasets after batch effect correction. (B) Corrected principal component analysis (PCA) diagram. (C-H) Heatmaps and volcano plots of DEGs from the three pairwise comparisons: (C, D) ATC vs. normal, (E, F) ATC vs. PTC, and (G, H) PTC vs. normal.

